# The quantification of protein-protein interaction interfaces using solvent-excluded surface-defined properties

**DOI:** 10.1101/294074

**Authors:** Lincong Wang

## Abstract

Protein-protein interaction (PPI) is the cornerstone of nearly every biological process. During last forty years PPI interfaces have been investigated extensively both *in vitro* and *in silico* in order to understand both the strength and specificity of PPI. At least three different models, the buried surface model, the O-ring model and the rim-and-core model, have been proposed for PPI interface. However none of them provide much detail about PPI and a single model that reconciles them remains elusive. To identify common physical and geometrical features shared by various PPI interfaces we have analyzed several solvent-excluded surface (SES)-defined properties for a set of well-studied protein-protein complexes with crystal structures. Our analysis shows that the SES-defined properties for the interface atoms of a PPI partner are in general different from those for the surface atoms of a water-soluble protein. Most significantly we find that the partially-buried atoms of a PPI partner have unique SES-defined properties that set them well apart from either the buried atoms or the accessible atoms. Based on distinct SES-defined properties for the accessible, buried and partially-buried atoms shared by various PPI interfaces we propose a new model specified by a list of SES-defined properties shared by various PPI interfaces. Our model is quantitative in nature and should be useful for PPI site identification, protein-protein docking and structure-based design of chemicals targeting PPI.

## 1 Introduction

Due to the fundamental roles played by protein-protein interaction (PPI^1^) in nearly all biological processes, the structures of PPI interfaces as well as the strength and specificity of PPI have been studied extensively using both experimental and computational methods [1, 2, 3]. The rational behind the structural characterization of PPI interfaces is the physical principle that the structure of a molecule determines its function. Much have been learned about PPI interface, binding affinity and to a less degree specificity. Though PPI interfaces are diverse [4, 5, 6] at least three models, the buried surface model [7, 8], the O-ring model [9] and the core-and-rim model [10], have been proposed to highlight their common features. The buried surface model arose from the analyses of the PPI interfaces in the crystal structures of protein-protein complexes. It defines an interface *geometrically* as a set of surface atoms^2^ whose solvent-accessible surface areas (ASAs) decrease upon binding. It states that every interface atom (or residue) contributes to binding affinity and its contribution is proportional to the amount of ASA that is buried upon complexation. A key piece of evidence for the buried surface model is the existence of a decent correlation between binding affinity and the total buried ASA of interface atoms or residues [7, 8, 11]. The O-ring model has its origin in alanine scanning mutagenesis experiment [12]. The model does not rely on structural information since it defines a PPI interface *operationally or thermodynamically*. In contrary to the buried surface model the O-ring model states that only a relatively small number of hot-spot residues contribute disproportionately to binding affinity. It agrees with the buried surface model in that most hot-spot residues have large buried ASAs, are near the center of an interface defined geometrically by the buried surface model and are hydrophobic in nature. The O-ring model states that the non-hot spot residues at an interface act to occlude bulk solvent from the hot-spots [9]. However the model provides no details about either the mechanism for solvent occlusion or how such occlusion contributes to PPI. As with the buried surface model the core-and-rim model defines an interface using the change in atomic ASA upon complexation. However unlike the former it goes one step further and uses atomic ASA to divide the interface atoms (residues) of a protein-protein complex into two different groups: a core and a rim. The core is composed mainly of hydrophobic residues while the rim is enriched in hydrophilic ones. At residue-level the core is generally more conserved than the rim [13]. The rim is accessible to bulk solvent. However the model provides no details about either protein-solvent interaction between rim atoms and solvent molecules or how the rim contributes to PPI. The O-ring model could account to some extent for binding specificity at residue-level while both the buried surface and core-and-rim models say little about it. None of the three models are specified using a quantitative relation between interface structure and binding affinity though they all agree that the inter-partner hydrophobic attraction ^3^ contributes largely to binding affinity. A quantitative model that reconciles them remains elusive at present. This illustrates a well-known challenge in structural analysis: the interpretation of thermodynamic data using structural information alone.

To identify common physical and geometrical properties shared by various PPI interfaces we have applied our accurate and robust solvent-excluded surface (SES) computation algorithm to a set of 143 well-studied protein-protein complexes with crystal structures [14]. In our analysis the surface atoms of a PPI partner^4^ *before* complexation are divided into three different sets: a set of solvent-accessible atoms and two sets of interface atoms: buried and partially-buried. Correspondingly the SES of a partner is divided into three regions: accessible, buried and partially-buried. Our buried and partially-buried regions are similar but not identical to the core and the rim^5^ of the core-and-rim model. Upon complexation the SES area of an accessible atom does not change, the area of a buried atom becomes zero while the area of a partially-buried atom decreases. In our analysis, a PPI interface is described in terms of SES-defined physical and geometrical properties. To the best of our knowledge no SES-defined quantities have ever been used on a large-scale to characterize PPI interfaces except for two reports [8, 4] where SES but not SES-defined physical properties have been applied to two small sets of protein-protein complexes.

Our analysis shows that the accessible atoms of a PPI partner have SES-defined properties almost identical to those for the surface atoms of a typical water-soluble protein [15]. In the contrary each of the two interface sets has its own unique SES-defined properties. As with the previous observations the buried region of an interface is enriched in apolar atoms^6^. Interestingly our analysis finds that compared with the SESs of water-soluble proteins the buried regions in the 143 complexes have, on average, positive net charge and 27% higher atom density (number of surface atoms per SES area). Most significantly the partially-buried region of a partner has rather unusual SES-defined properties that set it well apart from both the accessible and the buried regions. Specifically if we use as references the averages of the SES-defined properties for a large set of water-soluble proteins then those for the buried regions are on one side while those for the partially-buried regions the other. No previous PPI analyses have divided the surface atoms of a PPI partner into three different groups as we do. In addition they have focused mainly on the differences between the PPI interface of a PPI partner and the rest of its surface. Consequently previous analyses have failed to identify unique structural features for PPI interfaces [16, 17] that could set a PPI interface apart from the rest of protein surface. In fact, it is generally believed that to a large extent the interface residues (atoms) do not differ chemically from the accessible ones [16, 10]. Furthermore quantitative differences between *accessible* atoms (residues) and *partially-buried* atoms (residues) have hardly ever been reported. Instead one previous analysis claims that the rim of a PPI partner is very similar to the rest of its surface [10].

The large differences in SES-defined properties among the three subsets of surface atoms shared by all the PPI interfaces in the 143 complexes lead us to propose a new model for PPI interfaces, the interior-and-boundary model. In our model the set of buried atoms forms the interior of an interface while the set of partially-buried atoms establishes the boundary and acts to seal the interior from direct interaction with bulk solvent. The unique SES-defined properties shared by all the boundaries in the set of 143 complexes are consistent with them having evolved for sealing the interior as tightly as possible. The sealing is achieved mainly by the direct inter-partner hydrogen bondings between the two sets of partially-buried polar atoms in a PPI interface and the indirect inter-partner hydrogen bondings mediated by individual water molecules. The tight-sealing likely increases binding affinity through a reduction in energy penalty for a buried region to have an atom density well above the average for water-soluble proteins. The latter is unfavorable for protein-solvent interaction but likely increases the inter-partner VDW attraction between the two sets of buried atoms in a PPI interface. In addition the sealing likely increases the strength of inter-partner hydrogen bonding interaction involving a buried polar atom. The requirement for tight-sealing also contributes to binding specificity. To some extent our model is a refinement to the core-and-rim model at atomic level. In particular the boundary of our model is similar to the rim of the latter. However, our model differs from the core-and-rim model in important details. In our model both the interior and the boundary are specified quantitatively using SES-defined properties and their roles in PPI are explicitly defined. Compared with previous qualitative models for PPI interfaces the quantitative nature of our model likely makes it better suited for PPI site identification and prediction [18, 19], protein-protein docking [20] and structure-based design of chemicals targeting PPI [21].

## 2 Materials and Methods

### 2.1 The PPI data set and data preparation

A list of well-characterized and widely-used protein-protein complexes [14] is first downloaded online. A set of crystal structures for the listed complexes are then downloaded from the PDB. A set ℂ of 143 rigid-body complexes^7^ (section S1 of the Supplementary Materials) is used in our interface analysis. Each of the 143 complexes in ℂ is visualized and analyzed chain-by-chain using our molecular analysis and visualization program to extract its two PPI partners according to the specification in the downloaded list. There may exist more than one interface for a complex if the original PDB file includes more than one pair of partners. Protons are added using the program REDUCE [22] to any PDB structure that lacks their coordinates. Protonation likely affects the structural characterization of PPI interfaces since it changes surface property both geometrically and physically. For example more heavy atoms become buried or partially-buried with protonation. However a protonated protein is more realistic as far as physics is concerned.

The SES-defined properties for a set 𝕊 of 16, 483 water-soluble proteins with high quality non-redundant crystal structures are used as references for the quantification of PPI interfaces. As described previously [15] set 𝕊 excludes not only hyperthermophilic, anti-freeze, membrane and nucleic acid binding proteins but also any protein that has a bound compound with ≥ 20 heavy atoms. In addition the SES-defined properties for a set of 1, 314 extended conformations (set 𝕄_e_ [15]) are also used for comparison.

## 2.2 SES computation and the representation of PPI interfaces

The SESs and SES-defined geometrical and physical properties of a complex and its two PPI partners are computed as described in our companion papers [23, 15]. For easy exposition the two partners are denoted respectively as: **1** and **2**. The surface atoms of a partner are divided into three distinct sets: accessible **A**_P_, buried **B**_P_, and partially-buried **R**_P_ according to the changes in their SES areas upon complexation. Let *a*_*b*_(*i*) and *a*_*f*_(*i*) be respectively the SES areas of surface atom *i* of a partner before and after complexation. If *a*_*b*_(*i*) − *a*_*f*_(*i*) < *T*_*a*_ then atom *i* ∈ **A**_P_, else if *a*_*f*_(*i*) < *T*_*a*_ then *i* ∈ **B**_P_, and else *i* ∈ **R**_P_ where *T*_*a*_ is a threshold with a typical value of 0.01Å^2^. Correspondingly the SES of a partner is divided into three different surface regions: *A*_P_ composed of the SESs of the atoms in **A**_P_, *B*_P_ the SESs of the atoms in **B**_P_, and *R*_P_ the SESs of the atoms in **R**_P_. The set of interface atoms **I**_P_ includes both buried and partially-buried atoms, that is **I**_P_ = **B**_P_ ∪ **R**_P_, and the SES of a PPI interface is *I*_P_ where *I*_P_ = *B*_P_ ∪ *R*_P_. When it becomes necessary to distinguish the two partners their respective sets will be denoted explicitly as (1) and (2). For example **R**_P_(1) is the **R**_P_ of partner **1**, **R**_P_(2) the **R**_P_ of partner **2** and then **R**_P_ = **R**_P_(1) ∪ **R**_P_(2) is the set of partially-buried atoms for the complex.

## 2.3 The angular distribution of partially-buried polar atoms

The boundary (partially-buried region) of a typical PPI interface is enriched in polar atoms that are either hydrogen bond donors or acceptors [15]. The distribution of polar atoms along the boundary could be characterized as follows. Let **R**_P_(1) and **R**_P_(2) be respectively the sets of partially-buried atoms of partner **1** and **2** of a complex so **R**_P_ = **R**_P_(1) ∪ **R**_P_(2) is the set of the partially-buried atoms for its interface. The atoms in **R**_P_ are first fitted to a 2-dimensional (2D) plane 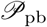 and their projections onto 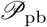 are then represented by 2D polar coordinates ((*r, θ*)s). Let 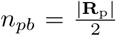 where |**R**_P_| is the number of atoms in **R**_P_. The fitted plane 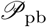 is then divided evenly into a list **u**_*a*_ of *n*_*pb*_ angular intervals with a unit polar angle 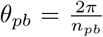 for each interval. The computation of the angular distribution for all the atoms in **R**_P_ proceeds as follows.

1. SET every element in **u**_*a*_ FALSE
2. MOVE the origin of the coordinate system to the center for all the atoms in **R**_P_
3. BEST-FIT all the atoms in **R**_P_ to a plane 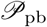 and let Δ_*pb*_ be the fitting residual
4. FOR each *polar* atom *i* in **R**_P_
5. PROJECT it onto 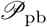
6. COMPUTE its 2D polar coordinate *r*_i_, *θ*_*i*_
7. COMPUTE its angular index 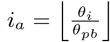
8. SET element (interval) *i*_*a*_ of **u**_*a*_ TRUE
9. COMPUTE average boundary radius 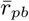 and angular coverage C_pb_

The average boundary radius 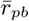 for an interface is defined as 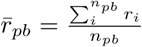 and its angular coverage C_pb_ defined as 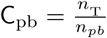 where *n*_T_ is the total number of TRUE intervals in **u**_*a*_.

## 2.4 The angular distribution of inter-partner hydrogen bonds between R_P_(1) and R_P_(2)

A polar atom in **R**_P_(1) could form an inter-partner hydrogen bond with a polar atom in **R**_P_(2) either directly or mediated by a water molecule. The distribution of such inter-partner hydrogen bonds between **R**_P_(1) and **R**_P_(2) along the boundary of a PPI interface could be computed as follows.

1. SET every element in **u**_*a*_ FALSE
2. FOR each polar atom *i* in **R**_P_(1)
3. FOR each polar atom *j* in **R**_P_(2)
4. IF both *i* and *j* are hydrogen atoms {a hydrogen-bond mediated by a water molecule}
5. IF *d*_*ij*_ < *T*_HH_
6. PROJECT *i, j* onto 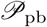
7. COMPUTE their polar angles *θ*_i_, θ_j_
8. COMPUTE their angular indexes 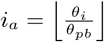 and 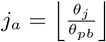
9. SET elements (intervals) *i*_*a*_ and *j*_*a*_ of **u***a* TRUE
10. ELSE IF either *i* or *j* is hydrogen atom AND *d*_*ij*_ < *T*_*XH*_ {either a direct or indirect hydrogen-bond}
11. PROJECT *i, j* onto 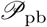
12. COMPUTE their polar angles *θ*_*i*_, *θ*_*j*_
13. COMPUTE their angular indexes 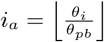 and 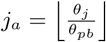
14. SET elements (intervals) *i*_*a*_ and *j*_*a*_ of **u**_*a*_ TRUE

The angular coverage *C*_hb_ for the polar atoms that could form an inter-partner hydrogen bond is defined as C_hb_ = 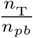 where *n*_T_ is the total number of TRUE intervals in **u**_*a*_. Thresholds *T*_HH_ and *T*_XH_ are used respectively to determine whether two polar hydrogen atoms could form a hydrogen bond mediated by a water molecule and whether a polar hydrogen atom could form a hydrogen bond with a polar heavy atom either directly or mediated by a water molecule. The value for *T*_HH_ is 4.0Å while *T*_*XH*_ is set to 3.5Å.

### 3 Results and Discussion

## 3.1 The SES-defined physical and geometrical properties of PPI interfaces

As described in our companion paper [15], we have identified a list of SES-defined physical and geometrical properties that are relevant to protein-solvent interaction. Among them are surface atom density (*ν*), the ratio of the number of surface apolar atoms over that of surface polar atoms (*n*^*oi*^), average-partial charge or charge per atom (*ρ*), the ratio of the total SES area of polar atoms over that of apolar atoms (*A*^*io*^), the ratio of SES area per apolar atom over that of per polar atom (*R*_*oi*_), and concave-convex ratio (*r*_*cc*_). In our definition a polar atom is either a hydrogen bond donor or acceptor while an apolar atom may not form a hydrogen bond with any other atoms [15]. For each PPI partner we compute these SES-defined properties for each of its three sets **A**_P_, **B**_P_ and **R**_P_. We expect that the SES-defined properties for the **I**_P_s in ℂ are to be different from those for 𝕊. For easy exposition the set of accessible atoms, the set of buried atoms and the SES of a water-soluble protein in 𝕊 are denoted respectively as **A**_S_, **B**_S_ and *A*_S_. The list of SES-defined properties especially their averages over the **A**_S_s in 𝕊 are used as references for the evaluation of the differences between the *I*_P_s in ℂ and the *A*_S_s in 𝕊. Specifically the differences in SES-defined properties between **B**_P_ and **A**_S_ and the differences between **R**_P_ and **A**_S_ are used to separate the *B*_P_ of a PPI interface from its *R*_P_ and to specify our interior-and-boundary model.

### 3.1.1 Surface atom density ***ν*** and the ratio of the number of surface polar atoms over that of surface apolar atoms *n*^*oi*^

A quantity that could be easily computed for a protein structure is its surface atom density 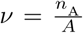 where *n*_*A*_ is the number of its surface atoms and *A* its total SES area. For a water-soluble protein with *n* < 10, 000 where *n* is its total number of atoms, *ν* increases slowly with *n* but remains on average the same when *n* > 10, 000 [15]. The average *ν* for all the **A**_S_s in 𝕊, 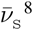, is 0.0938 atom per Å^2^. The average *ν*s for the **A**_P_s, **B**_P_s and **R**_P_s in ℂ are respectively 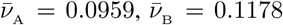 and 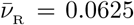 atoms per Å^2^ (Fig. 1a). It shows that in terms of surface atom density, the **A**_P_s are almost identical to the **A**_S_s while 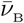 is 27% higher but 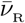 25% lower than 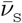. As with the *ν*s the *n*^*oi*^s also have different values for the three sets of surface atoms (Fig. 1b). As to be expected the average *n*^*oi*^ for the **A**_P_s is 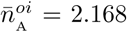, close to the average 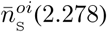 for 𝕊 while the averages for the **B**_P_s and **R**_P_s are respectively 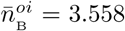 and 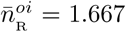, the former is about 56.2% *larger* while the latter is about 26.8% *smaller* than 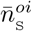. Thus compared with the SES of a water-soluble protein the buried region of a PPI interface not only has higher atom density but also more apolar atoms. That the **B**_P_s have high atom densities is consistent with the previous observation that PPI interfaces are generally tightly-packed [7, 24]. A higher *ν* and a larger *n*^*oi*^ in combination with the shape complementary between **B**_P_(1) and **B**_P_(2) [25] lead to stronger VDW attraction between them. However a *ν* value and a *n*^*oi*^ value both close to their corresponding averages for 𝕊 are preferred for optimal interaction with bulk aqueous solvent. Thus the benefit of having a high *ν* and a large *n*^*oi*^ for any surface region could only be realized if that region is sealed from direct interaction with bulk solvent.

**Figure 1:**
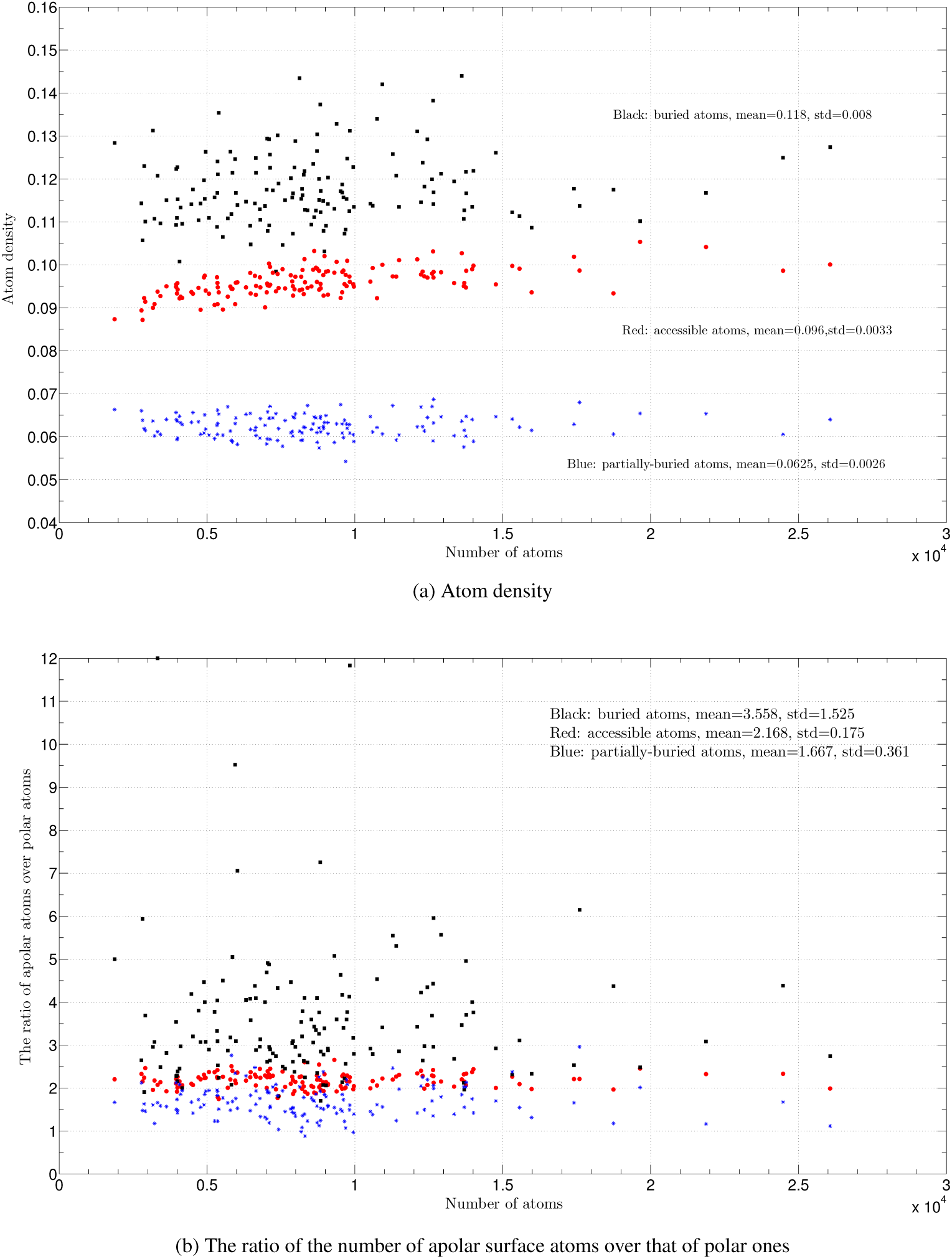
The *ρ*s and the *n*^*oi*^s for the A_P_s, B_P_s and R_P_s in ℂ. In both (**a**) and (**b**) the sets of buried, accessible and partially-buried atoms are depicted respectively as filled squares, filled circles and stars and are colored respectively in black, red and blue. The inserts in both (**a**) and (**b**) list their respective means and standard deviations. For references the average *ρ*s and the average *n*^*oi*^ for 𝕊 are respectively 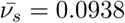 atom per Å^2^ and 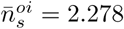. The x-axis is the number of atoms in a protein-protein complex. The y-axis is atom density *ρ*s in figure (**a**) and ratio *n*^*oi*^ in figure (**b**).

In contrast with the 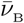 and 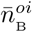, the 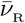 is lower and 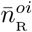 (Fig. 1) smaller than their corresponding averages for 𝕊. If we assume that the **A**_S_s in 𝕊 have evolved for optimal interaction with bulk solvent, then a *ν* lower and a *n*^*oi*^ smaller than their corresponding averages for 𝕊 mean that the partially-buried region of a PPI interface must have evolved for something more than just optimal interaction with bulk solvent. As to be detailed later a key function of the two partially-buried regions of a PPI interface is to use their abundant inter-partner hydrogen bonds to seal its buried region (interior) from direct interaction with bulk solvent.

Interestingly the *ν*s in ℂ satisfy the following inequalities: *ν*_B_ > *ν*_A_ > *ν*_R_ (Fig. 1a). Except for three complexes (1e96, 1fqj and 1oph) the *n*^*oi*^s for all the other ones in ℂ satisfy the following inequality: 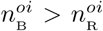, and except for nine complexes the *n*^*oi*^s for all the other ones satisfy the following inequality: 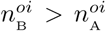. It means that all the *ν*s and the great majority of the *n*^*oi*^s for the **B**_P_s are well separated from those for either the **A**_P_s or the **R**_P_s. In addition compared with the *ν*s and *n*^*oi*^s for both the **A**_P_s and the **R**_P_s, the *ν*s and *n*^*oi*^s for the **B**_P_s have much larger ranges (Fig. 1). These large variations among individual interfaces reflect PPI diversity [4, 5, 6, 14] in ℂ and suggest the relevance of their individual values to binding affinity and specificity. In summary the SES-defined properties *ν* and *n*^*oi*^ are likely useful for PPI site detection, protein-protein docking and the design of chemicals targeting PPI.

### 3.1.2 The surface charge of PPI interface

The water-soluble proteins in 𝕊 all have *negative* net surface charges with an average 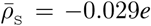 per surface atom while the **B**_S_s in 𝕊 all have *positive* net charges with an average 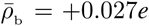 per buried atom. As shown in Fig. 2 the *ρ*s for the **B**_P_s in ℂ have a 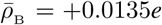, that is, the *ρ*s of the individual sets of buried atoms in ℂ are on average *positive*. If we equate hydrophobicity with *ρ*, we conclude that the **B**_P_s are, on average, only half as hydrophobic as the interiors of the water-soluble proteins in 𝕊. The latter likely helps the dissolution into aqueous solvent of the individual uncomplexed partners in ℂ [14]. However, according to previous studies [26, 27] it is unlikely that the enrichment of positive charges in **B**_P_ contributes directly to binding affinity.

**Figure 2:**
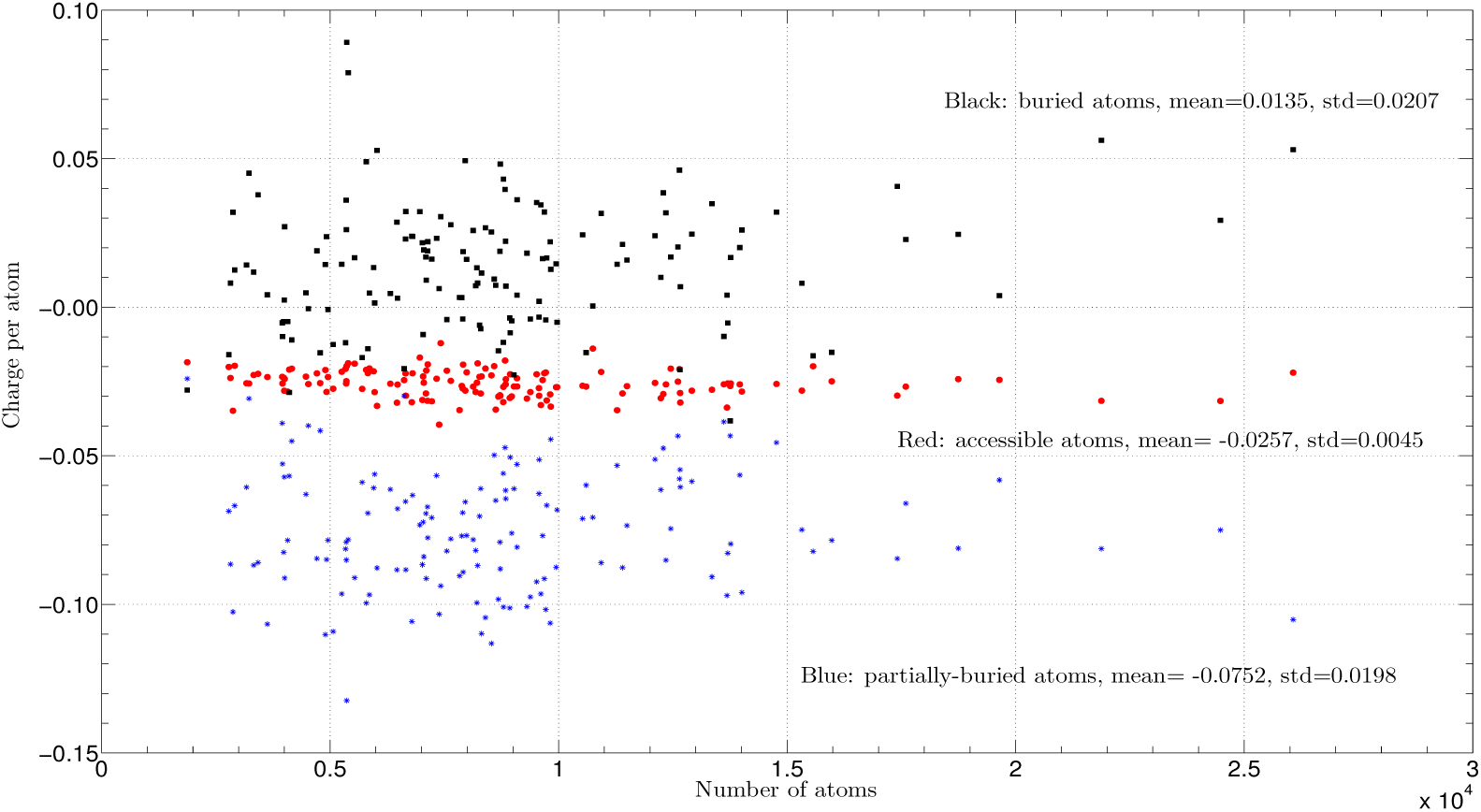
The ρs for the A_P_s, B_P_s and R_P_s in ℂ. The three inserts list their respective means (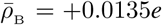, 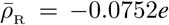 and 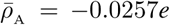 per atom) and standard deviations. For comparison the average *ρ* for all the accessible atoms in 𝕊 is 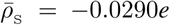 per atom. The x-axis is the number of atoms in a protein-protein complex. The y-axis is charge per atom in *e*.

In stark contrast to the **B**_P_s in ℂ, the average *ρ* for the **R**_P_s is 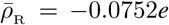 per atom, that is a 2.6-fold more *negative* than 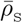. As shown in Fig. 6c a negative *ρ* value for a **R**_P_ is due mainly to the presence of hydrogen donors (oxygen and nitrogen atoms) in the **R**_P_. As described previously, a 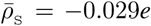 per atom is optimal for protein-solvent interaction [15]. Thus on average close to 61.4% _9_ of the polar atoms in the **R**_P_s must have evolved for something else. According to previous studies [26, 27] it is unlikely that the enrichment in the **R**_P_s of the atoms with negative charges increases binding affinity. On the other hand, such enrichment likely increases the number of inter-partner hydrogen bonds between **R**_P_(1) and **R**_P_(2). As to be detailed later the polar atoms in a **R**_P_ distribute rather evenly along the boundary of a PPI interface and the inter-partner hydrogen bonds between **R**_P_(1) and **R**_P_(2) cover close to half of the boundary. Taken together the results suggest that the polar atoms in a boundary help seal the set of buried atoms (the interior) from direct interaction with bulk solvent by means of many inter-partner hydrogen bonds between **R**_P_(1) and **R**_P_(2).

Interestingly in terms of *ρ* the three sets of surface atoms are well separated from each other (Fig. 2). In fact, all the PPI interfaces in ℂ satisfy *ρ*_R_ < *ρ*_A_ and 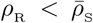. Only a single complex, 2oob [28] (the complex of the UBA domain of Cbl-b and ubiquitin), has 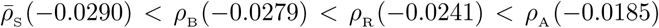. This complex with a total of 1,876 atoms is the smallest among the 143 ones in ℂ. Though the hydrophobic interaction between UBA and ubiquitin has been described in much detail in the original paper and is thought to be important for PPI [28], this interface is in fact the most hydrophilic one among all the interfaces in ℂ as judged by *ρ*_B_ < *ρ*_R_ < *ρ*_A_. The existence of a possible dimerization interface between two UBAs [28] may explain why this particular complex has the least negative *ρ*_A_ value among all the complexes in ℂ^10^. Only three complexes, 1fle, 2oob and 4fqi, have their *ρ*_*A*_s more positive than their *ρ*_B_s 𝕊. In addition compared with the *ρ*s for either **A**_P_ or **A**_S_, the *ρ*s for both **B**_P_ and **R**_P_ have much larger ranges (Fig. 2). Their large variations reflect the diversity of the PPIs in ℂ and suggest their relevance to binding affinity and specificity. In summary our analysis shows that the SES-defined *ρ* is relevant to PPI and thus is likely useful for PPI site detection, protein-protein docking and the design of chemicals targeting PPI.

### 3.1.3 The ratio of the SES area of apolar atoms over that of polar atoms

The ratio of the total SES area of the apolar atoms over that of the polar atoms of a water-soluble protein, 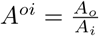 has been shown to be relevant to protein solubility [15]. The average *A*^*oi*^ for 𝕊 is 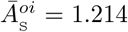. As to be expected the *A*^*oi*^ average for the **A**_P_s in ℂ is 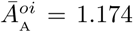, rather close to 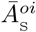. However, the average *A*^*oi*^ for the **B**_P_s is 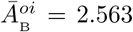, a 2.11-fold *increase* over 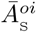. A more than 2-fold increase in the total SES area of buried apolar atoms likely enhances the inter-partner hydrophobic attraction between **B**_P_(1) and **B**_P_(2). Very interestingly the average *A*^*oi*^ for the **R**_P_s is only 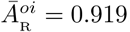, less than 1.0 and a 24.3% *decrease* over 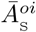. The former means that the polar surface area_11_ of a partially-buried region is larger than its apolar surface area while the latter is consistent with the conclusion that the polar atoms in a partially-buried region participate in inter-partner hydrogen bonding. Significantly as with *ν* and *n*^*oi*^, the *A*^*oi*^s for the **B**_P_s have a large range with a standard deviation of 1.613 (Fig. 3). In contrary the standard deviation for the 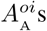 is only 0.109 and that for the 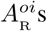 is only 0.226. It suggests that in terms of *A*^*oi*^ the PPIs in ℂ are highly varied in their buried regions and thus *A*^*oi*^ is likely pertinent to binding affinity and specificity. In addition, out of the 143 complexes in ℂ only two complexes, 1ahw 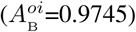 and 3mxw 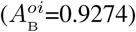, have 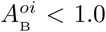. The former [29] is an antigen-antibody complex with the antigen being an extracellular domain while the latter [30] is an antigen-antibody complex with the antigen being a secreted protein. Having 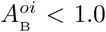 may reflect the uniqueness of their PPI interfaces. Only three complexes, 1e96, 1oph and 2j0t, have 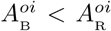 and only four complexes, 1ahw, 1e96, 2gtp and 3mxw, have 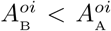. Thus the SES-defined property *A*^*oi*^ is likely useful for PPI site detection, protein-protein docking and the design of chemicals targeting PPI.

**Figure 3:**
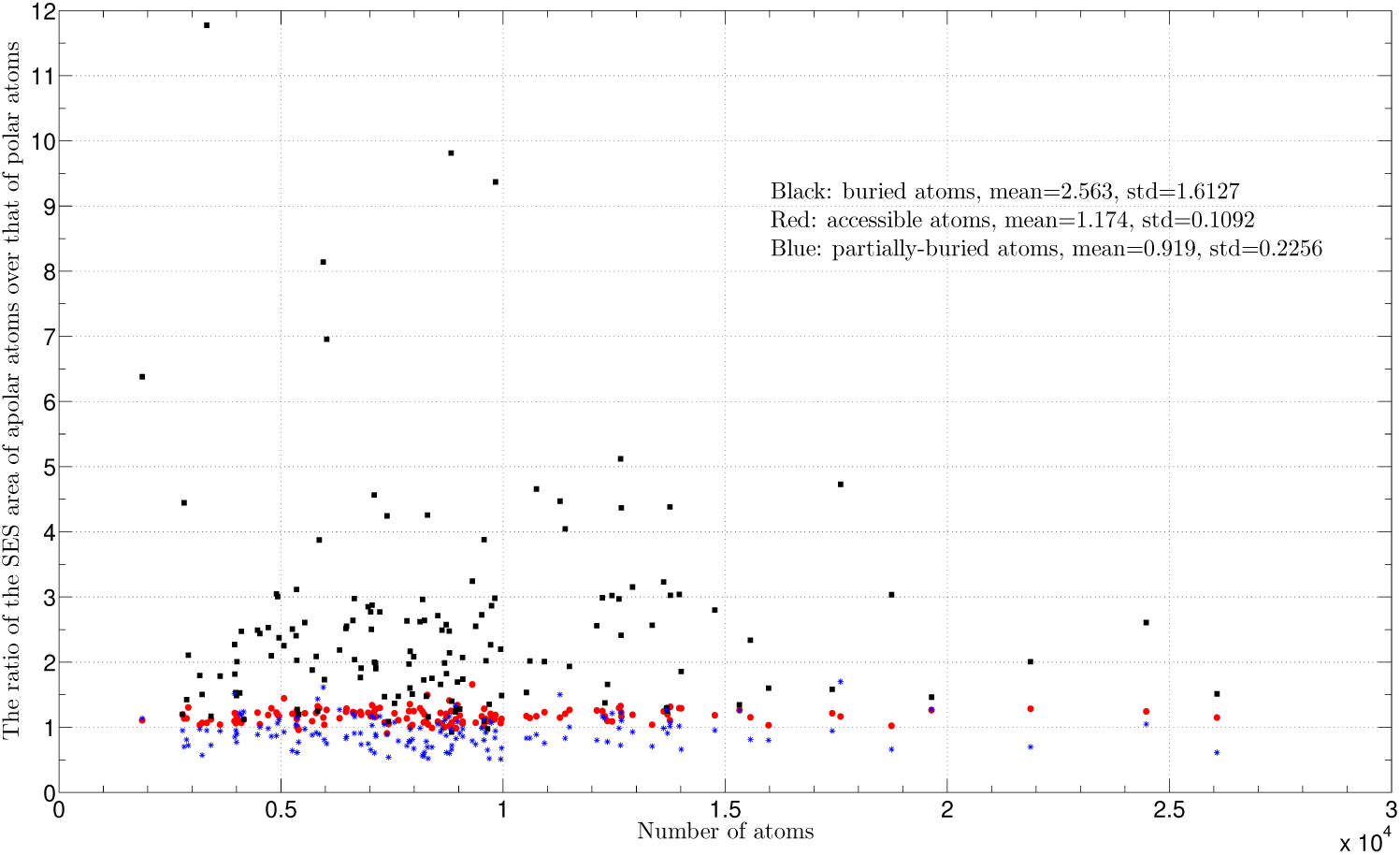
The *A*^*oi*^s for the A_P_s, B_P_s and R_P_s in ℂ. The insert lists their respective means, 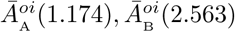 and 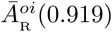, and standard deviations. For comparison the average *A*^*oi*^ for the **A**_S_s in 𝕊 is 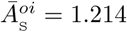 and out of 16, 483 water-soluble proteins in 𝕊 only 92 (0.56%) proteins have 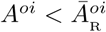. The x-axis is the number of atoms in a protein-protein complex. The y-axis is *A*^*oi*^.

**Figure 4:**
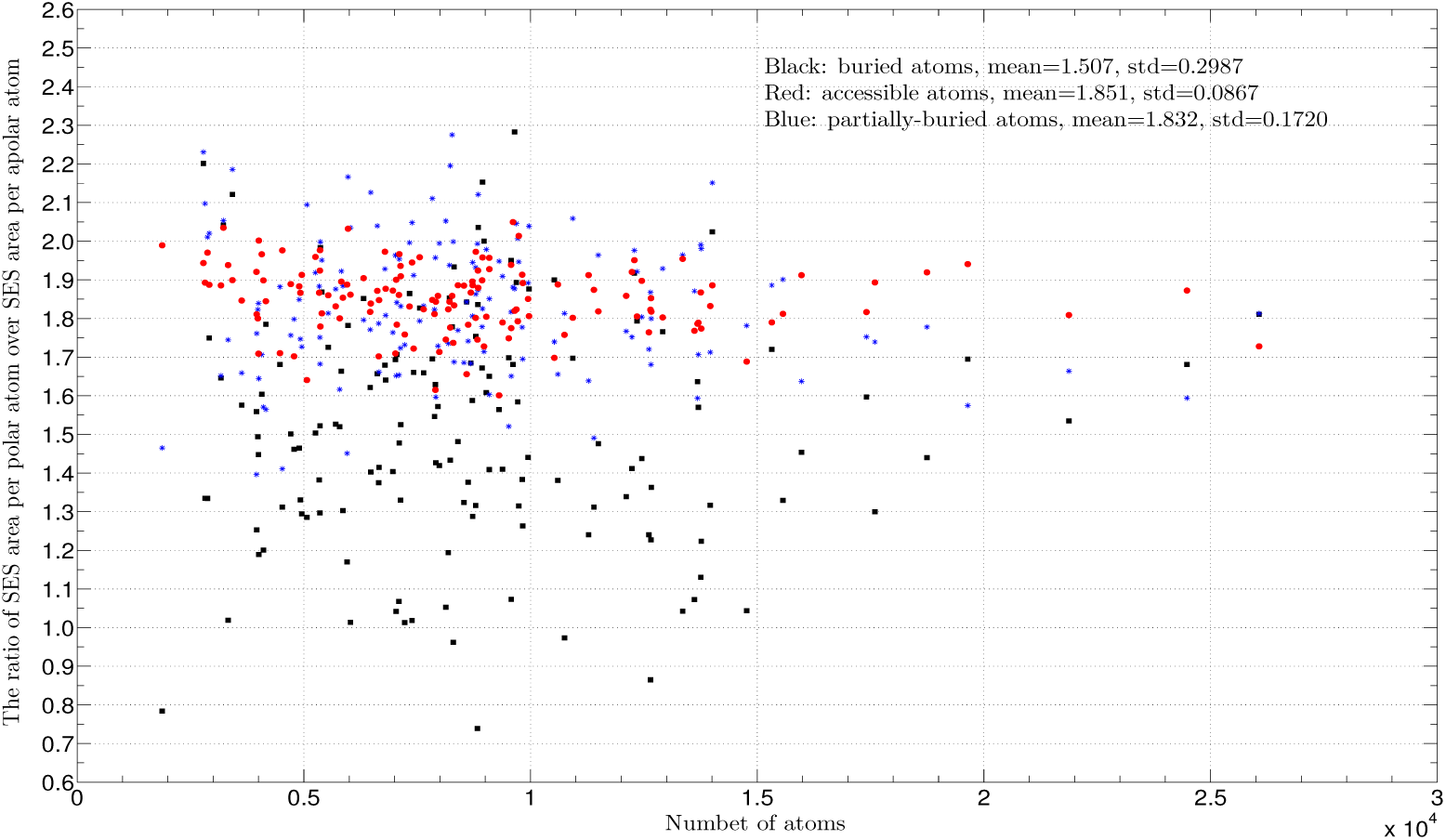
The *R*^*io*^s for the A_P_s, B_P_s and R_P_s in 𝕊. The insert lists their respective means 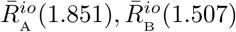 and 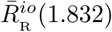 and standard deviations. For comparison, the average *R*^*io*^ for the **A**_S_s in 𝕊 is 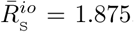. The x-axis is the number of atoms in a protein-protein complex. The y-axis is *R*^*io*^.

The *R*^*io*^s for 𝕊 have a 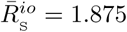. We hypothesize that the increase in SES area per polar atom optimizes the intermolecular hydrogen bonding between the surface polar atoms of a water-soluble protein and solvent molecules. As to be expected the average *R*^*io*^ for the **A**_P_s, 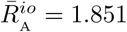, is rather close to 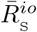. However the average *R*^*io*^ for the **B**_P_s is only 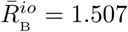. There are only five proteins, 2ouw, 2qsk, 2rfr, 3qva and 4z0m, out of 16, 483 proteins in 𝕊 that have their *R*^*io*^s slightly smaller than 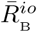 [15]. Thus compared with the SESs of water-soluble proteins, the **B**_P_s are, on average, not only positively-charged, enriched in apolar atoms but their polar atoms also have smaller SES area per atom. This reduction in SES area for a buried polar atom could be easily visualized as shown in Fig. 6a. Since a *B*_P_ region is sealed from direct interaction with bulk solvent, there is no need to optimize intermolecular hydrogen bonding between surface polar atoms and solvent molecules. On the other hand the reduced SES area for a polar atom may affect the strength of its hydrogen bonding interaction in subtle way. The reduction in SES area likely leads to sub-optimal inter-partner hydrogen bonding interaction between **I**_P_(1) and **I**_P_(2) [31] but the apolar nature of a *B*_P_ region could possibly compensate for it by increasing the strength of such hydrogen bonding interaction. In addition since the uncomplexed partners in ℂ are soluble themselves, the reduced area per polar atom may decrease the hydrogen bonding interaction with bulk solvent molecules and consequently makes them less soluble. The average *R*^*io*^ for the **R**_P_s is 1.832, slightly smaller than 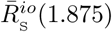 and 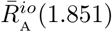 (1.851) and about 21.6% larger than 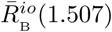. It suggests that the polar atoms in the **R**_P_s are almost as capable as those in the **A**_S_s to form the intermolecular hydrogen bonds between themselves and with solvent molecules.

### 3.1.4 The concave-convex ratios of interface atoms

The concave-convex ratio 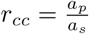 of a surface atom is a SES-defined geometrical property that quantifies its local smoothness. A SES region with a small *r*_*cc*_ is locally-rugged and likely tightly-packed. Typically such a region has more carbon atoms than a region with a larger *r*_*cc*_. As described in our companion paper [15] the *r*_*cc*_s for the surface apolar atoms of a water-soluble protein in 𝕊 are, on average, 1.5-fold larger than the *r*_*cc*_s for the surface polar atoms, and the *r*_*cc*_s for extended conformations are several-fold smaller than those for folded structures. In addition there exist good correlations between surface residue *r*_*cc*_s and several known hydrophobicity scales [15]. We hypothesize that from a protein-solvent interaction perspective a locally-smooth region composed mainly of apolar atoms is better than a locally-rugged region with similar composition. Compared with the latter the former is less disruptive to water’s hydrogen-bonded network and has stronger VDW attraction with solvent molecules.

As shown in Fig. 5 and Table 1 of section 3.3, the *r*_*cc*_s for either the polar or apolar atoms in the **A**_P_s are close to those in the **A**_S_s. However the 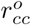 average for the **B**_P_s is smaller than that for either the **A**_P_s or the **A**_S_s in 𝕊. This decrease in 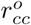 is consistent with both the abundance of apolar carbon atoms in a *B*_P_ and the preference of aromatic hydrophobic residues in a PPI interface. In the contrary the average 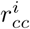 for the **B**_P_s is slightly larger than that for the **A**_P_s. It suggests that even a buried polar atom is somewhat optimized for inter-partner VDW attraction. Furthermore the 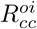 average for the **B**_P_s is only 1.267, smaller than the average 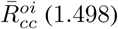 (1.498) for 𝕊 (section S2 and Figure S1 of the Supplementary Materials). The local ruggedness of a buried region as judged by the smallness of its *r*_*cc*_ does not contradict its being globally flat (or more properly shallow) as reported in previous PPI interface analyses [16, 21]. A small *r*_*cc*_, a high *ν* and a large *n*^*oi*^ for a *B*_P_ as well as its global shallowness are all consistent with that the buried atoms especially the buried apolar atoms are tightly packed against each other [7]. In combination with the shape complementary between *B*_P_(1) and *B*_P_(2) the tight packing possibly maximizes the inter-partner VDW attraction between them and thus contributes largely to binding affinity and specificity.

**Table 1:** The averages of eight SES-defined properties for six sets of surface atoms. The eight averages are respectively 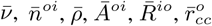 (apolar atoms), 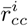 (polar atoms) and 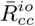. The six sets of surface atoms are respectively **B**_P_, **R**_P_, **I**_P_, **A**_P_, **A**_S_ and the set of surface atoms in 𝕄_e_ [15]. Except for 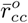 and 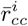 the other six averages over 𝕄_e_ lie between those for the **B**_P_s and those for the **R**_P_s. Except for 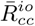, the other seven averages for the **I**_P_s in ℂ lie between those for the **B**_P_s and those for the **R**_P_s.

| | $\bar{\nu}$ | $\bar{n}^{oi}$ | $\bar{\rho}$ | $\bar{A}^{oi}$ | $\bar{R}^{io}$ | $\bar{r}_{cc}^i$ (polar) | $\bar{r}_{cc}^o$ (apolar) | $\bar{R}_{cc}^{oi}$ |
| --- | --- | --- | --- | --- | --- | --- | --- | --- |
| $\mathbf{B}_p$ | 0.1178 | 3.558 | +0.0135 | 2.563 | 1.506 | 0.289 | 0.339 | 1.267 |
| $\mathbf{R}_p$ | 0.0625 | 1.667 | -0.0752 | 0.919 | 1.832 | 0.175 | 0.231 | 1.343 |
| $\mathbf{I}_p$ | 0.0790 | 2.216 | -0.0359 | 1.189 | 1.705 | 0.195 | 0.270 | 1.414 |
| $\mathbf{A}_p$ | 0.0959 | 2.184 | -0.0257 | 1.174 | 1.851 | 0.283 | 0.414 | 1.477 |
| $\mathbf{A}_s$ | **0.0938** | **2.278** | **-0.0290** | **1.241** | **1.875** | **0.267** | **0.392** | **1.496** |
| $\mathbb{M}_e$ | **0.0706** | **2.457** | **+0.00037** | **1.570** | **1.567** | **0.119** | **0.139** | **1.31** |

**Figure 5:**
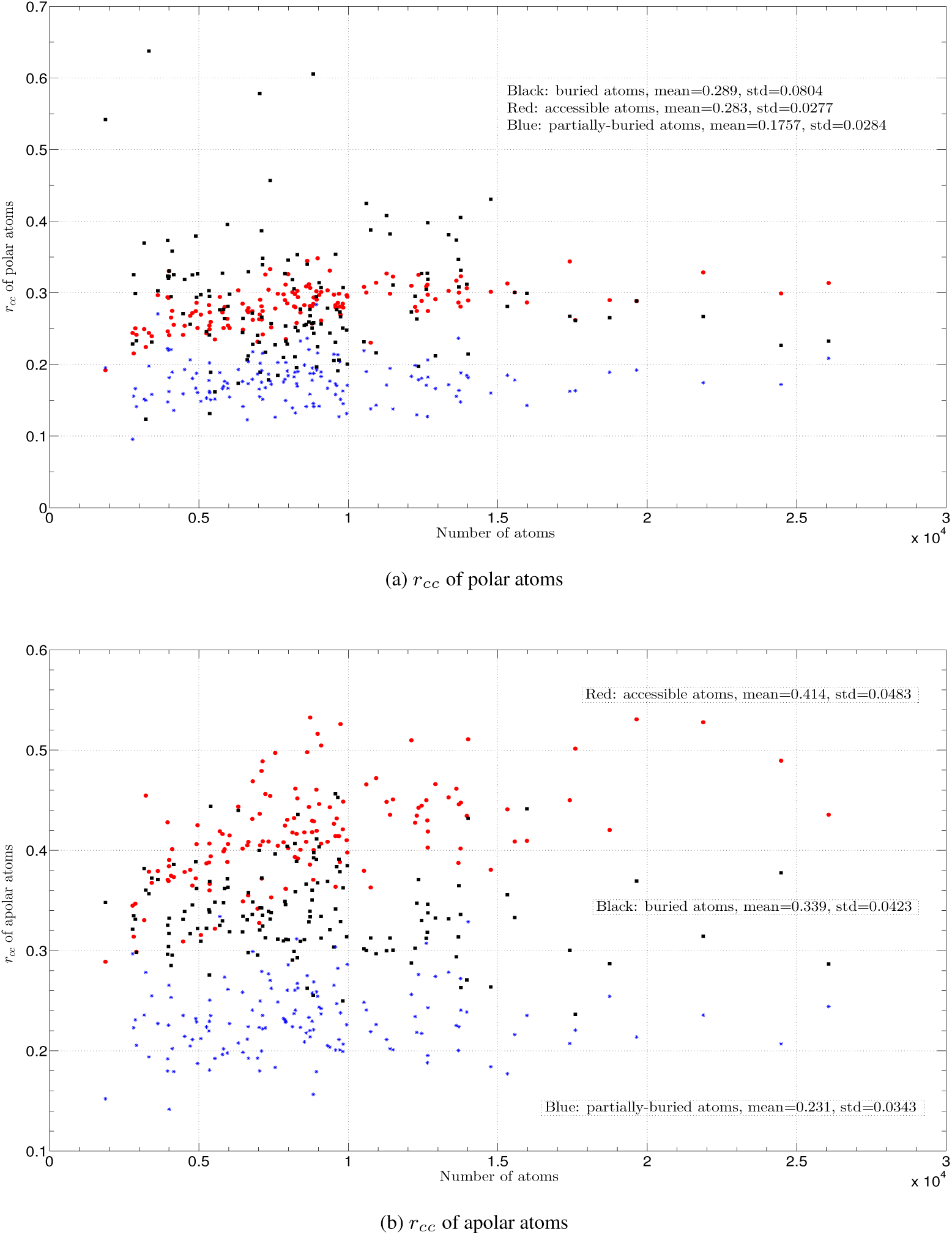
The concave-convex ratios (*r*_*cc*_s) of the polar atoms and apolar atoms of the *A*_P_s, *B*_P_s and *R*_P_s in ℂ. Figures **a** and **b** show respectively the *r*_*cc*_s for the polar and apolar atoms. The insert in (**a**) and the three inserts in (**b**) list their respective means and standard deviations. The x-axis is the number of atoms in a protein-protein complex. The y-axis is *r*_*cc*_.

Surprisingly the *r*_*cc*_ average for the **R**_P_s is *smaller* than that for the **B**_P_s. Particularly the average *r*_*cc*_ for the apolar atoms in the **R**_P_s is only 0.231 (Fig. 5b), somewhat close to 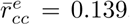, the *r*_*cc*_ average for the apolar atoms in M_*e*_ [15]. Furthermore no PPI partner in ℂ has a *r*_*cc*_ value for **R**_P_ larger than that for **A**_P_. Compared with a surface region with a large *r*_*cc*_, a surface region with a small *r*_*cc*_ has either more of tightly-packed atoms or less carbon atoms or both. As described in section 3.1.1 compared with either the **A**_P_s or the **A**_S_s in 𝕊, the **R**_P_s have lower *ν*s and smaller *n*^*oi*^s. The atoms in a surface region with a lower *ν* have less neighbors while a surface region with a small *n*^*oi*^ has relatively abundant surface polar atoms. The fact that the three averages (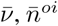 and 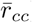) are all small is consistent with the abundance of hydrogen bond donors such as oxygen and nitrogen atoms in the **R**_P_s. From a protein-solvent interaction perspective this fact shows that the **R**_P_s must have evolved for more than optimal interaction with bulk solvent molecules. The latter is likely achieved through optimal intermolecular hydrogen bonding and VDW attraction between protein surface atoms and solvent molecules. Based on these unique SES-defined properties for the **R**_P_s we conclude that a partially-buried region has evolved for optimal inter-partner hydrogen bonding interaction between **R**_P_(1) and **R**_P_(2). Such inter-partner hydrogen bonds make it possible for a **R**_P_ to occlude the buried region from direct interaction with bulk solvent.

As with other SES-defined properties the *r*_*cc*_ range for the **B**_P_s is larger than that for the **A**_P_s (Fig. 5a). In addition for each interface in ℂ and for both the polar and apolar atoms the *r*_*cc*_ values for the **B**_P_s are larger than those for the **A**_P_s. There are only two complexes, 1a2k and 1ahw, that have the 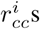 for their **R**_P_s larger than those for their **B**_P_s. There are also only two complexes, 1avx and 1bvn, that have the 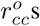 for their **R**_P_s larger than those for their **B**_P_s. Taken together our analysis shows that SES-defined property *r*_*cc*_ is likely useful for PPI site detection, protein-protein docking and the design of chemicals targeting PPI.

## 3.2 The angular distribution of partially-buried polar atoms

As described above compared with both the **A**_P_s and the **A**_S_s the **R**_P_s have lower atom density, more polar atoms, larger negative partial charge per atom, smaller *A*^*oi*^ and smaller *r*_*cc*_. Based on these characteristic SES-defined properties we conclude that the partially-buried atoms have evolved not only for optimal interaction with bulk solvent molecules but more likely for the formation of inter-partner hydrogen bonds either directly or indirectly via individual water molecules. If the conclusion is correct, we expect that the polar atoms in a **R**_P_ distribute more or less evenly along the boundary. As with Kolmogorov-Smirnov test we first divide a fitted 2D polar plane into 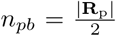 angular intervals, we then compute angular coverage 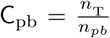 for each *R*_P_ to determine the distribution uniformity of its polar atoms. As shown in Fig. 6, the average angular coverage *C*_pb_ over the individual sets of the polar atoms in the **R**_P_s is 0.798. It means that on average an interface in ℂ has only 20% of its boundary *not* covered by any polar atom that could potentially form a hydrogen bond with either protein atoms or solvent molecules. The abundance of individual water molecules along the rims of PPI interfaces has been reported in many crystal structures and has been visually illustrated before [4]. Since the partially-buried polar atoms in a **R**_P_ have, on average, large negative partial charges, their near even distribution along the boundary likely reduces the energy penalty caused by the electrostatic repulsion among themselves. If we select only the polar atoms that could form an inter-partner hydrogen bond either directly or indirectly via a water molecule, on average these polar atoms still cover, on average, 38.2% of a boundary (Fig. 6). Interestingly the *C*_hb_ range is relatively large with a standard deviation of 0.108 and a mean of 0.382. The large variation in individual *C*_hb_s suggests that the *C*_hb_ distribution along a PPI boundary may pertain to binding specificity.

**Figure 6:**
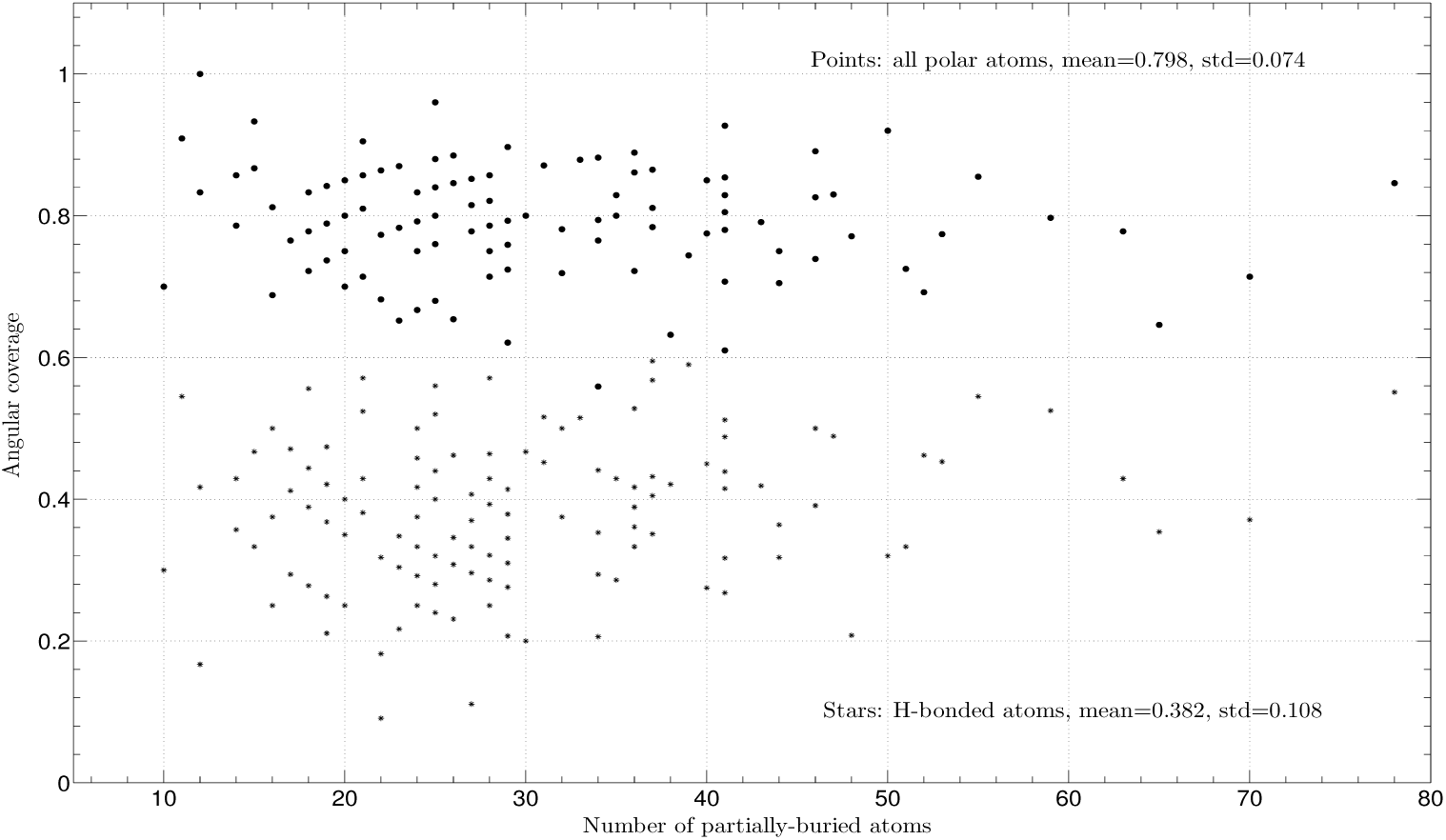
The angular coverage of the individual sets of the polar atoms in the R_P_s. The two inserts list the respective means and standard deviations for the *C*_pb_s and the *C*_hb_s in ℂ. The *C*_pb_s are depicted as points and the *C*_hb_s as stars. The x-axis is the number of partially-buried atoms in an interface, |**R**_P_|, in a complex. The y-axis is angular coverage.

The angular distribution is computed by best-fitting all the atoms in a **R**_P_ to a 2D plane. A fitting residual **R**_pb_ for a **R**_P_ quantifies its planarity. As shown in Fig. 7, the change of Δ_pb_ with the number of the partially-buried atoms in an interface, |**R**_P_|, is slightly linear with a slope of 0.056, an intercept of 2.659Å and a *R*_square_ = 0.259. The smallness of the slope shows that Δ_pb_ increases very slowly with |**R**_P_|. To obtain an estimation for the geometrical size of a *R*_P_ we first move the origin of the coordinate system to the center for all its atoms and then compute the polar radius *r*_pb_ for each partially-buried atom and finally compute their average boundary radius 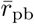. The change of 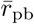 with |**R**_P_| is modestly linear with a slope of 0.089 and a *R*_square_ = 0.454 (Fig. 7). This fitted linear equation between 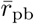 and |**R**_P_| may be useful for PPI site identification and protein-protein docking.

**Figure 7:**
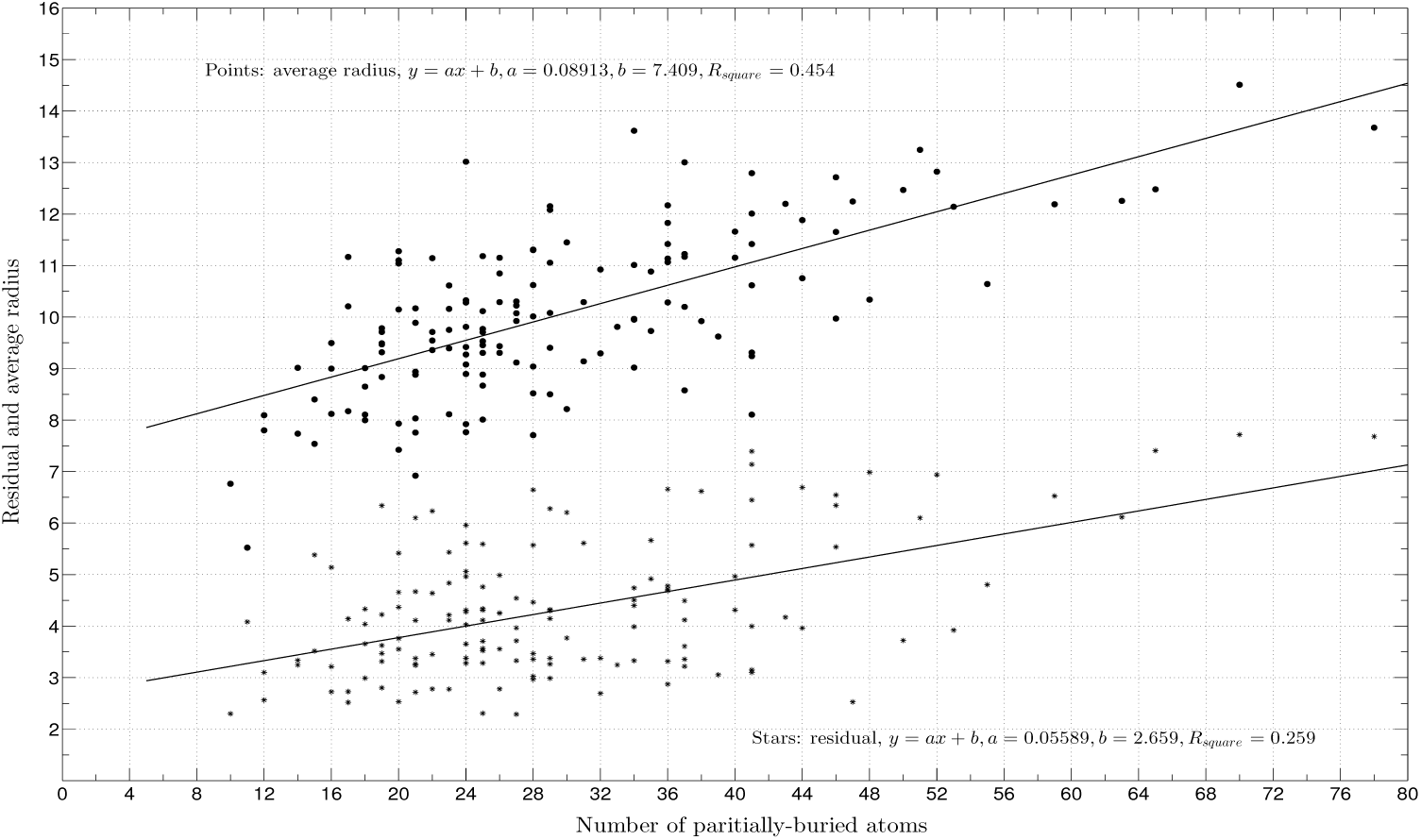
The fitting residuals and average boundary radii of the R_P_**s in** C. Each residual (Δ_pb_) of best-fitting all the atoms in an individual **R**_P_ to a 2D plane is depicted as a point. The average radii 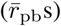 are depicted as stars. Both residual Δ_*pb*_ and 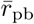 increase somewhat linearly with the number of the partially-buried atoms (|**R**_P_|) with a *R*_square_ = 0.259 for the former and a *R*_square_ = 0.454 for the latter. The x-axis is |**R**_P_|. The y-axis is either residual or average boundary radius with a unit of Å.

## 3.3 The interior-and-boundary model for PPI interfaces

In our analysis the surface atoms of a PPI partner is divided into three different sets: a set of solvent-accessible atoms (**A**_P_) and two sets of interface atoms: a set of buried atoms (**B**_P_) and a set of partially-buried atoms (**R**_P_). As described above both the **B**_P_s and the **R**_P_s in ℂ have their own characteristic SES-defined physical and geometrical properties that are different from those for a large set of water-soluble proteins (set 𝕊). Most interestingly if we use as references the averages for a list of eight SES-defined properties for 𝕊, then most of the SES-defined properties for the **B**_P_s are on one side while those for the **R**_P_s the other (Table 1). More specifically the 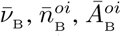 and 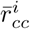 for the buried atoms are all larger than those for 𝕊 while 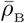 is positive (Table 1). In stark contrast the 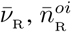, and 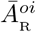 and 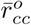 for the partially-buried atoms are all smaller than those for 𝕊 while 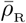 is 2.6-fold more negative than 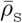. However if we first merge a **B**_P_ and a **R**_P_ into a single set **I**_P_, and then compute the averages over all the **I**_P_s in ℂ for the list of SES-defined properties, the differences between the averages for the **I**_P_s and those for either the **A**_P_s or 𝕊 are rather small (Table 1). Interestingly the SES-defined properties for the **I**_P_s are somewhat close to those for 𝕄_*e*_. In contrast with our division of the surface atoms of a PPI partner into three different sets, the majority of the previous structural analyses on PPI interfaces did not separate them into different groups. Instead they focus mainly on the differences between the **I**_P_s and the **A**_P_s. Though the separation of an interface into a core and a rim [32, 10] is similar to our division the difference between the rim and the core and the difference between the rim and the rest of protein surface (set **A**_P_) have not been quantified in the past using any surface-defined properties. In particular the core-and-rim model provides no details for either protein-solvent interaction between rim atoms and solvent molecules or how the rim contributes to PPI. In fact it has been claimed that the rim of a PPI interface is very similar to the rest of protein surface [10]. The previous structural analyses have so far failed to identify common geometrical or physical properties shared by various interfaces that could be used to separate a PPI interface from the rest of protein surface [16, 17]. Consequently both the buried surface model and the rim-and-core model are qualitative in nature and do not provide much detail about PPI interfaces. Specifically both have much difficulty to interpret the alanine scanning mutagenesis data. On the other hand the O-ring model that does not rely on structural information assigns no roles to a large percentage of the residues in a PPI interface. In addition the model provides no details for either the mechanism for solvent occlusion or how such occlusion contributes to PPI. Up to now a single model that reconciles the differences among the three models remains elusive.

Based on the quantitative differences in SES-defined properties among the **A**_P_s, **B**_P_s and **R**_P_s in ℂ and by comparing their averages with those for 𝕊, we propose the interior-and-boundary model for PPI interfaces with the following common features. Each PPI interface is composed of an interior (a buried region) and a boundary (partially-buried region). The interior is enriched in apolar atoms, tightly-packed, locally-rugged and has positive net charge. Upon complexation it is sealed by the boundary from direct interaction with bulk solvent molecules. The interior of a PPI partner has likely evolved for optimal VDW interaction with the buried atoms of its partner rather than for optimal VDW or intermolecular hydrogen bonding interaction with bulk solvent molecules. The interior thus makes disproportionately large contribution to PPI affinity. The variations in SES-defined properties for the **B**_P_s are much larger than those for the **A**_P_s and **R**_P_s. It means that the interior also determines binding specificity. The boundary is enriched in polar atoms, loosely-packed, locally-smooth and has large negative net charge. The polar atoms in the boundary distribute evenly along the boundary and the polar atoms that are capable of forming hydrogen bonds cover about 40% of the boundary. The boundary has evolved to a large degree for optimal inter-partner hydrogen bonding interaction and to a less degree for optimal interaction with bulk solvent molecules, and acts to seal the interior from direct interaction with bulk solvent. The occlusion of bulk solvent from the buried region of partner **1** enhances its attraction to the buried region of partner **2** through the inter-partner VDW and hydrogen bonding interactions, and thus contributes indirectly to binding affinity. The variations in SES-defined properties for the **R**_P_s are larger than those for the **A**_P_s. The boundary thus contributes to PPI affinity and specificity as a whole though the contribution by each individual residue or atom is relatively small. The O-ring model does state that the non-hot spot residues at an interface act to occlude bulk solvent from its hot-spots [9]. However the non-hot spot residues in the O-ring model may not be in the boundary specified in our model.

The large differences between an interior and a boundary could be easily visualized using SESs. As shown in Fig. 8a the differences in surface area between the partially-buried atoms and buried atoms and between the accessible atoms and buried atoms are less pronounced when a VDW surface is used: the sole buried oxygen atom and the sole buried nitrogen atom each has a large surface area in the VDW surface but each has a much smaller spherical polygon area *a*_*s*_. The buried polar atoms are much less exposed on a SES than on a VDW surface. Thus in terms of molecular visualization a SES is more sensitive than a VDW surface in showing the geometrical and physical properties that are relevant to both protein-solvent interaction and PPI. SESs have been used before for the visualization of PPI interfaces in order to illustrate their diversity rather that for the SES’s relevance to PPI [4].

**Figure 8:**
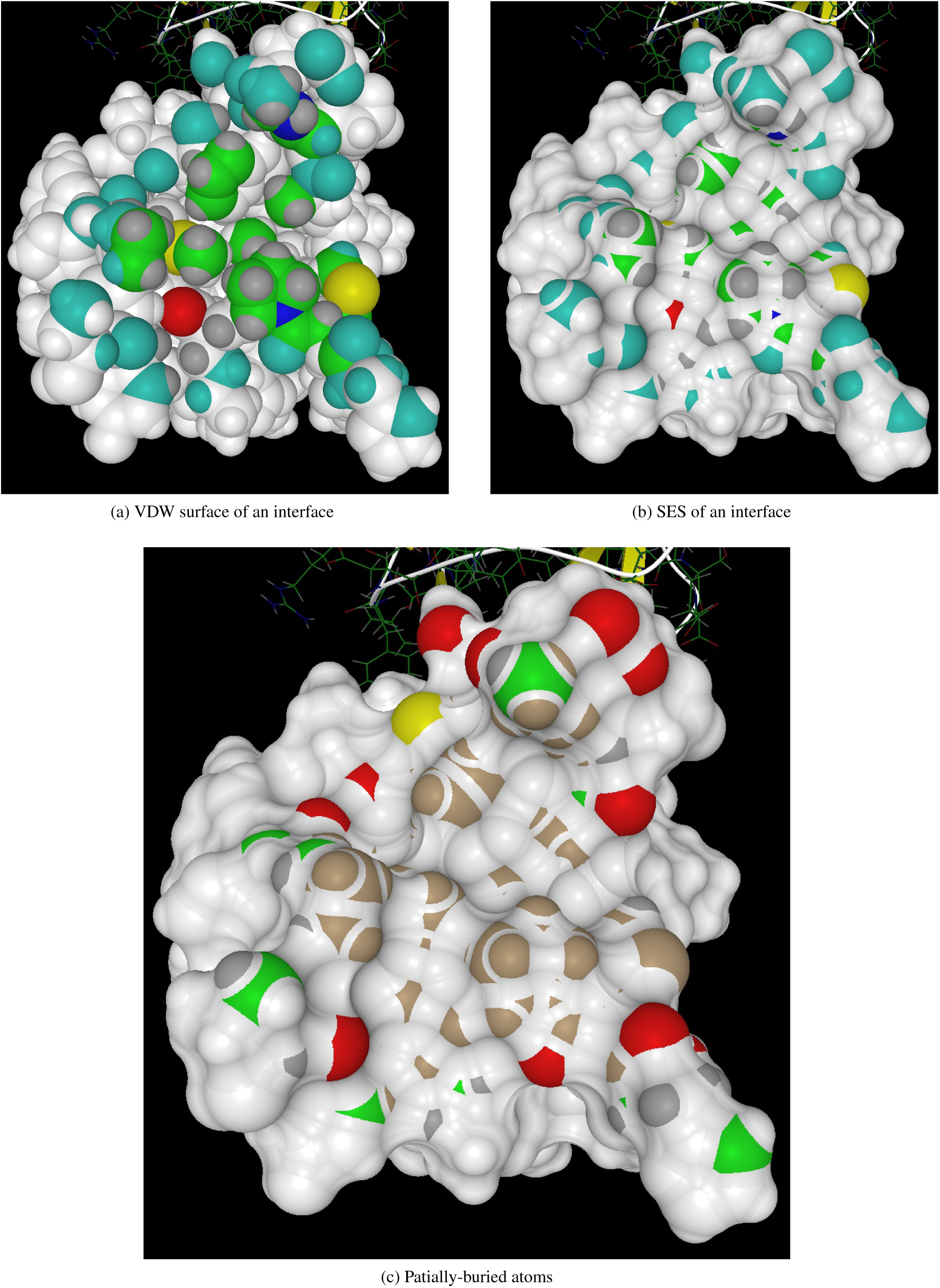
The buried and partially-buried atoms of a typical PPI partner. All the partially-buried atoms in the upper two figures are colored in cyan while buried C, O, N, 𝕊 and H atoms are colored respectively in green, red, blue, yellow and dark gray. In figure (**c**) all the buried atoms are colored in brown while the partially-buried C, O, N, 𝕊 and H atoms are colored respectively in green, red, blue, yellow and dark gray. The three figures are prepared using our structural analysis and visualization program written in C++/Qt/OpenGL.

## 3.4 The potential applications to PPI site identification, protein-protein docking and the design of compounds again PPI

Unlike the previous three models for PPI interfaces, our interior-and-boundary model is quantitative in nature with key features specified by a list of SES-defined properties. These characteristic SES-defined properties shared by all the analyzed PPI interfaces should be useful for PPI interface identification given the structural information for a partner. Similarly they could be used for protein-protein docking. Due to the fundamental roles played by PPI in nearly all biological processes PPI interfaces have become key targets for therapeutic drugs especially small molecule compounds. However it is still very challenging to design small compounds against PPI due at least in part to the lack of understanding of detailed physical and geometrical properties of PPI interfaces. Our discovery of a list of distinct SES-defined properties shared by various PPI interfaces should be useful for guiding the design of compounds with SES-defined properties that match those in a PPI interface.

### 4 Conclusion

Protein-protein interactions (PPIs) underlie nearly all biological processes. Though the structures of a large number of protein-protein complexes are currently available, the quantitative relationship between the structure of a PPI interface on the one hand and PPI strength and specificity on the other hand is largely unknown. To better understand the roles of interface atoms in PPI we have analyzed a set of well-studied PPI interfaces using solvent-excluded surface (SES) and SES-defined physical and geometrical properties. The results show that in terms of SES-defined properties the boundary (the set of partially-buried atoms) of a PPI interface differs significantly from its interior (the set of buried atoms), and both the boundary and interior differ largely from the solvent-accessible atoms of a water-soluble protein. Specifically the unique SES-defined properties shared by the boundaries of the analyzed PPI interfaces suggest that the boundaries have evolved mainly for the sealing of the interiors by means of abundant inter-partner hydrogen bonds. The characteristic SES-defined properties for the interiors suggest that they have evolved mainly for inter-partner VDW attractions. From the SES-based analysis we propose the interior-and-boundary model for PPI interfaces. The model is specified in terms of quantitative SES-defined properties shared by various PPI interfaces and thus should be useful for PPI site identification, protein-protein docking and the design of chemicals targeting PPI.

## Supplementary Materials

### S1 The list of protein-protein complexes

Our analysis of PPI interfaces is performed on 143 rigid-body complexes whose pdbids are: 1a2k, 1ahw, 1akj, 1avx, 1ay7, 1azs, 1bj1, 1buh, 1bvk, 1bvn, 1clv, 1d6r, 1dfj, 1dqj, 1e6e, 1e6j, 1e96, 1efn, 1ewy, 1exb, 1f34, 1f51, 1fcc, 1ffw, 1fle, 1fqj, 1fsk, 1gcq, 1ghq, 1gl1, 1gla, 1gpw, 1gxd, 1h9d, 1hcf, 1he1, 1hia, 1i4d, 1i9r, 1iqd, 1j2j, 1jps, 1jtd, 1jtg, 1jwh, 1k4c, 1k74, 1kac, 1klu, 1ktz, 1kxp, 1kxq, 1m27, 1mah, 1ml0, 1mlc, 1nca, 1nsn, 1oc0, 1ofu, 1oph, 1oyv, 1ppe, 1pvh, 1qa9, 1qfw, 1r0r, 1rlb, 1rv6, 1s1q, 1sbb, 1t6b, 1tmq, 1udi, 1us7, 1vfb, 1wdw, 1wej, 1xd3, 1xu1, 1yvb, 1z0k, 1z5y, 1zhh, 1zhi, 2a1a, 2a5t, 2a9k, 2abz, 2ajf, 2ayo, 2b42, 2b4j, 2btf, 2fd6, 2fju, 2g77, 2gaf, 2gtp, 2hle, 2hqs, 2i25, 2j0t, 2jel, 2mta, 2o8v, 2oob, 2oor, 2oul, 2pcc, 2sic, 2sni, 2uuy, 2vdb, 2vis, 2vxt, 2w9e, 2x9a, 2yvj, 3a4s, 3biw, 3bp8, 3d5s, 3eoa, 3h2v, 3hmx, 3k75, 3lvk, 3mxw, 3p57, 3pc8, 3rvw, 3sgq, 3vlb, 4cpa, 4dn4, 4fqi, 4g6j, 4g6m, 4h03, 4hx3, 4m76, 7cei.

### S2 The 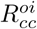 of PPI interface

As described in the main text the ratio of the concave-convex ratio for the apolar atoms 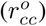 of a water-soluble protein over the concave-convex ratio for its polar atoms 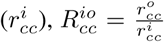, likely pertains to protein-solvent interaction. As shown in Figure S1, the 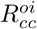 average for the buried atoms is 15.3% smaller than that for 𝕊 while 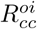 average for the partially-buried atoms is only 10.0% smaller that that for 𝕊. The 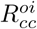 average for the **A**_P_s is rather close to that for 𝕊. As with other SES-defined properties, the 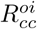 average for the **I**_P_s is close to that for the **A**_P_s.

**Figure S1:**
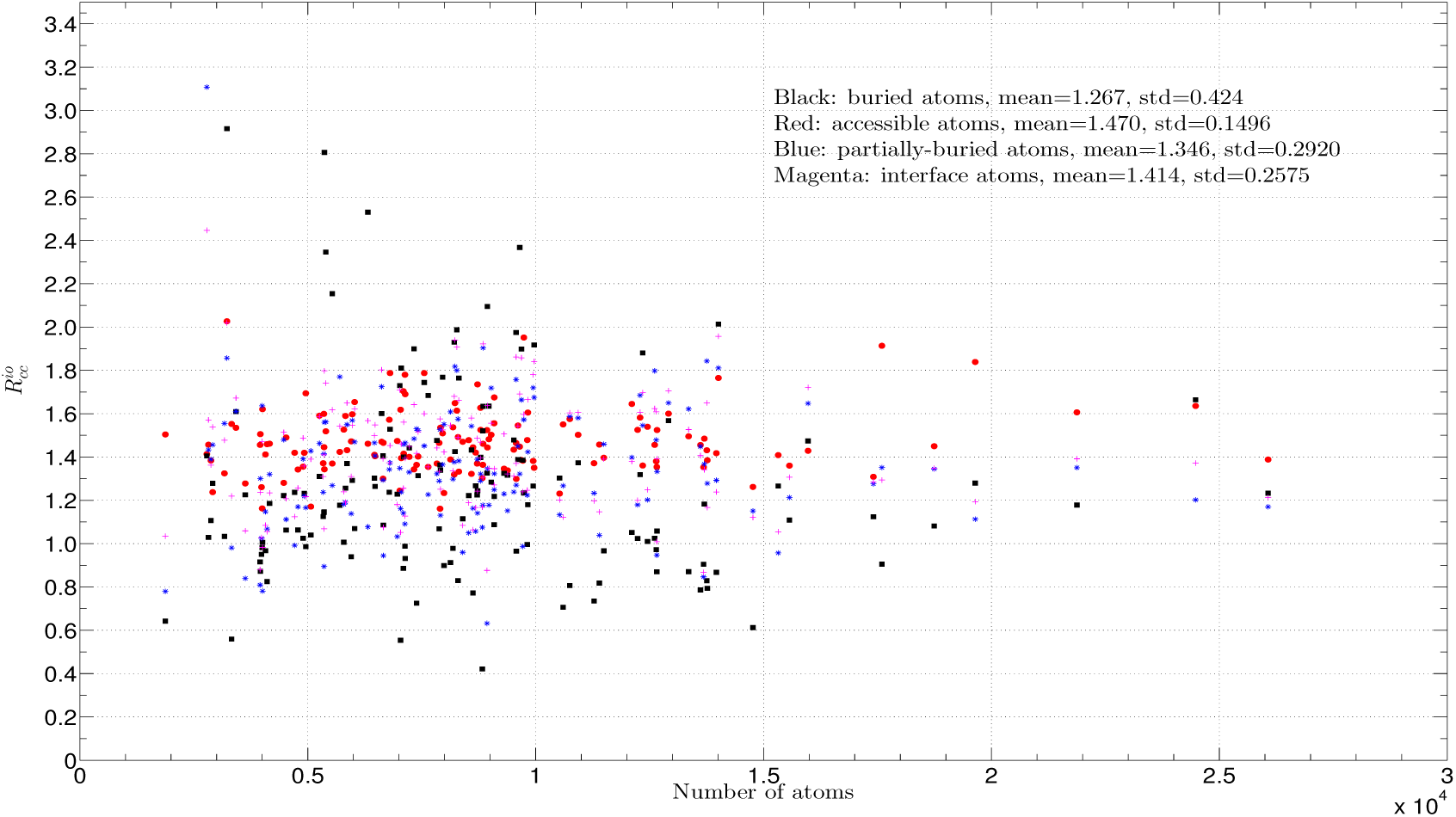
The 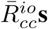 for the A_P_s, B_P_s, R_P_s and I_P_s in ℂ. The 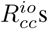 for the **A**_P_s, **B**_P_s, **R**_P_s and **I**_P_s are colored respectively in red, black, blue and magenta. The insert lists their respective means, 1.470, 1.267, 1.346 and 1.414, and standard deviations. For comparison the average 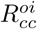 for the **A**_S_s in 𝕊 is 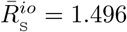. The x-axis is the number of atoms in a protein-protein complex. The y-axis is 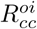.

## Footnotes

1 Abbreviations: SES, solvent-excluded surface; SAS, solvent-accessible surface; VDW, van der Waals; PPI, protein-protein interaction; 2D, two-dimensional; PDB, Protein Data Bank.

2 In this paper the term *surface* atom means either an atom in the set of all the solvent-accessible atoms of a PPI partner before complexation or an atom in the set of all the solvent-accessible atoms of a water-soluble protein. The two terms *solvent-accessible* atom and *accessible* atom are reserved mainly for the surface atoms whose SES areas do not change after complexation.

3 The term hydrophobic attraction or interaction is qualitative in nature when being used to describe PPI in solution since a hydrophobic residue or an apolar atom at a PPI interface is almost always surrounded by some hydrophilic residues or polar atoms. The latter makes it very difficult to quantify the hydrophobic attraction in PPI either experimentally or computationally.

4 A PPI partner may be composed of more than one chain while an interface is always formed between two partners: partner **1** and **2**. Each partner has its own set of interface atoms (residues). An interaction between **1** and **2** will be described as *inter-partner* interaction.

5 For brevity *rim* and *partially-buried region (boundary)* may be used interchangeably at qualitative level when there exists no ambiguity even though they are not the same.

6 An *apolar* atom may not form a hydrogen bond with other atoms while a *polar* atom is either a hydrogen bond donor or acceptor.

7 They are called rigid-body cases in Table BM5.xlsx. The original list, Table BM5.xlsx, includes 151 rigid-body cases. Eight complexes are excluded in our analysis due to some quality issues in the crystal structures as detected by our molecular analysis and visualization program.

8 A SES-defined property *x* for a set **A**_S_ in 𝕊 is denoted as *x*_S_. The average *x*_S_ over all the **A**_S_s is denoted as 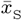. Similarly the *x*s for the three sets **A**_P_, **B**_P_ and **R**_P_ of a PPI partner are denoted respectively as *x*_A_, *x*_B_ and *x*_R_ and their averages over ℂ as 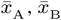, and 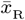. For brevity the average *x* over all the **A**_S_s in 𝕊 will be written as the average x for 𝕊.

9 The percentage is computed as follows: 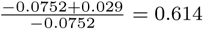

10 In our analysis this possible dimerization interface belongs to the accessible region (*A*_P_) of the complex.

11 Please see our companion paper [15] for the definitions of *polar surface area* and *apolar surface area*.

